# Benchmarking of proximity-dependent biotinylation enzymes across cellular compartments and time windows

**DOI:** 10.1101/2025.08.29.668156

**Authors:** Saya Sedighi, Kosar Vafaee, William Rod Hardy, Vesal Kasmaeifar, Zhen-Yuan Lin, Rawan Kalloush, Julia Kitaygorodsky, Brendon Seale, Queenie Hu, Monica Hasegan, Anne-Claude Gingras

## Abstract

Proximity-dependent biotinylation has become a powerful approach for mapping protein interactions and subcellular organization in living cells. Although a growing number of engineered biotin ligases have been introduced, their performance has not been systematically evaluated across diverse cellular contexts. Here, we benchmark ten proximity ligases spanning three bacterial lineages using standardized proteomic workflows across multiple labeling durations, subcellular compartments, and two human cell types. While all enzymes efficiently detect proximal associations, they differ in labeling kinetics, background activity, and spatial specificity. TurboID exhibits the highest overall activity but generates substantial background in standard media. miniTurbo and ultraID support rapid, biotin-dependent labeling with low background, making them better suited for dynamic and time-resolved applications. However, miniTurbo showed aberrant mitochondrial localization with two cytoskeletal baits (VASP and PFN1). Across 15 diverse baits, ultraID consistently provides an excellent combination of specificity, efficiency, and spatial compatibility—including unique recovery of Golgi-resident glycosyltransferases. This study serves as a comparative resource, offering guidance for enzyme selection and experimental design in proximity proteomics.

## Introduction

Understanding how proteins are organized within living cells is essential for uncovering the mechanisms that govern cellular function. Over the past decade, proximity-dependent biotinylation has emerged as a powerful approach to map protein neighbourhoods in live cells, organisms, and even fixed tissues ^1, 2^. These methods work through the covalent attachment of biotin to proteins in close proximity to a tagged “bait” protein, most commonly with engineered biotin ligases (e.g., BirA* and its derivatives) or peroxidases (e.g., APEX, derived from ascorbate peroxidase)^3–5^. The covalent nature of biotinylation enables stringent cell lysis conditions that disrupt protein complexes, membranes, chromatin, and the cytoskeleton. Biotin-labeled proteins can then be isolated by streptavidin affinity purification and identified by mass spectrometry^6^. When combined with well-matched negative controls and robust statistical analysis, proximity labeling thus enables the identification of true proximal interactors over background contaminants^7, 8^. A landmark study introduced BioID (Biotin Identification) to map protein associations of lamin A (*LMNA*) in HEK293 cells^6^. BioID is based on a mutant version of the *E. coli* biotin ligase BirA (R118G), commonly referred to as BirA*^6, 9–11^. In its wild-type form, BirA transfers biotin to carboxylases via a two-step mechanism: first generating biotinyl-AMP, which is retained through hydrogen bonding with the R118 side chain, and then transferring it to a lysine on the biotin carboxyl carrier protein (BCCP). The R118G mutation disrupts this retention, allowing biotinyl-AMP to diffuse away and react with lysines on nearby proteins, enabling proximity-dependent labeling^9–13^.

Since its introduction, BioID has had a major impact on the study of protein interactions and subcellular organization, enabling the detection of weak associations and the analysis of otherwise insoluble components^14–22^. However, the first-generation BirA* enzyme has relatively low catalytic activity. While shorter labeling times (1–3 h) have been reported^15, 23^, most experiments have typically used 12–24 h of labeling to achieve sufficient signal-to-noise levels to differentiate from background signals, limiting its utility for studying dynamic processes that occur within shorter timeframes. Peroxidase-based methods such as APEX offer minute-scale labeling but rely on hydrogen peroxide and phenol-based substrates, restricting their use *in vivo* or in settings sensitive to oxidative stress. To overcome the limitations of BirA*, several next-generation biotin ligases have been developed, using orthologs from other bacterial species^24–27^, domain deletions^24, 26^, point mutations^24, 25, 27^, reconstruction of ancestral variants^28^, and directed evolution^26, 29^.

Biotin ligases fall into three structural classes that all share a catalytic and C-terminal domain. Class I ligases, such as *Aquifex aeolicus* BirA, consist only of these domains, while Class II ligases (e.g., from *E. coli* and *Bacillus subtilis*) include an additional N-terminal DNA-binding domain that regulates the biotin biosynthesis operon^7, 11^. Targeted mutagenesis of the Class II *B. subtilis* ligase—via N-terminal deletion and R124G (analogous to R118G in *E. coli* BirA*), E323S, and G325R substitutions—yielded BASU, reported to show enhanced labeling kinetics^27^. Consistent with the Class I ligase structure, *A. aeolicus* BirA with the analogous R40G mutation^25^ generated BioID2, a smaller (∼26.4 kDa) enzyme than *E. coli* and *B. subtilis* BirA* (∼36 kDa), which, however, showed limited improvements in labeling efficiency^30–32^. Further structure-guided deletions of BioID2 at its C-terminus by two different groups, along with targeted point mutations, resulted in microID and microID2, compact variants (∼20 kDa) that retain catalytic activity and support faster labeling^24, 26^. Optimization of microID2 through targeted mutagenesis produced low-background microID2 (lbmicroID2), which exhibits reduced background^26^. Ancestral reconstruction of biotin ligases produced AirID, which shares ∼80% sequence identity with *E. coli* BirA*^28^ and is reported to have faster labeling kinetics, although it still operates on the timescale of hours.

A major leap in the development of biotin ligase-based proximity labeling tools was achieved through yeast-display-based directed evolution of *E. coli* BirA mutants. TurboID (∼35 kDa) and miniTurbo (∼27 kDa) were derived from an *E. coli* R118S background with 14 and 12 additional mutations, respectively, along with an N-terminal deletion in miniTurbo^29^. Both enzymes—particularly TurboID— exhibit high labeling efficiency, enabling robust biotinylation within minutes in cell culture^29^. This efficiency has supported widespread use of TurboID across diverse organisms, including *Drosophila*^33^, zebrafish^34, 35^, and yeast^36^. These variants have also enabled time-resolved proximity labeling^37–41^. In parallel, directed evolution of microID variants led to the development of ultraID (∼19 kDa). UltraID is a double mutant (R40G/L41P) that achieves TurboID-like kinetics with efficient labeling in as little as 10 min, while maintaining low background^26^.

While several studies have benchmarked individual or subsets of biotin ligase variants^24, 26, 28, 42^, none have systematically compared all enzymes side by side in complete mass spectrometry-based workflows. Here, we benchmarked ten published biotin ligases, and selected the top-performing ones for further analysis across different time points, diverse subcellular compartments, and two cell lines. Using LMNA as a reference bait, we evaluate both steady-state (6 h; all enzymes) and short labeling windows (5–60 min; selected enzymes) to reflect the growing interest in capturing dynamic changes in protein proximity. We then assess the top-performing ligases from each bacterial lineage using six additional baits spanning diverse subcellular compartments in HEK293 cells. Finally, we evaluate the top two enzymes with eight additional baits in HeLa cells. While all ligases efficiently recover proximal partners, they differ markedly in labeling kinetics, baseline activity, and compartmental specificity. TurboID generates the highest signal but exhibits high background in standard media, whereas miniTurbo and ultraID support rapid, biotin-dependent labeling with low background—making them better suited for time-resolved or inducible applications. Across the 15 baits tested, ultraID consistently combines low background with robust proximity labeling across diverse compartments, supporting its use as a preferred enzyme for studies requiring short labeling times and minimal constitutive activity.

## Results

### Generation of an expression toolbox for ten biotin ligases

We first generated a Gateway-compatible expression plasmid toolbox that supports the fusion of bait proteins to 10 biotin ligases from three bacterial species (Fig. 1a). Each vector includes a FLAG epitope tag and allows for either N- or C-terminal tagging (see Methods and Supplementary Table 1). The vector backbone, pcDNA5, enables inducible expression by tetracycline (or doxycycline), and targeted integration at a single genomic locus in Flp-In T-REx cell systems.

**Fig. 1.**
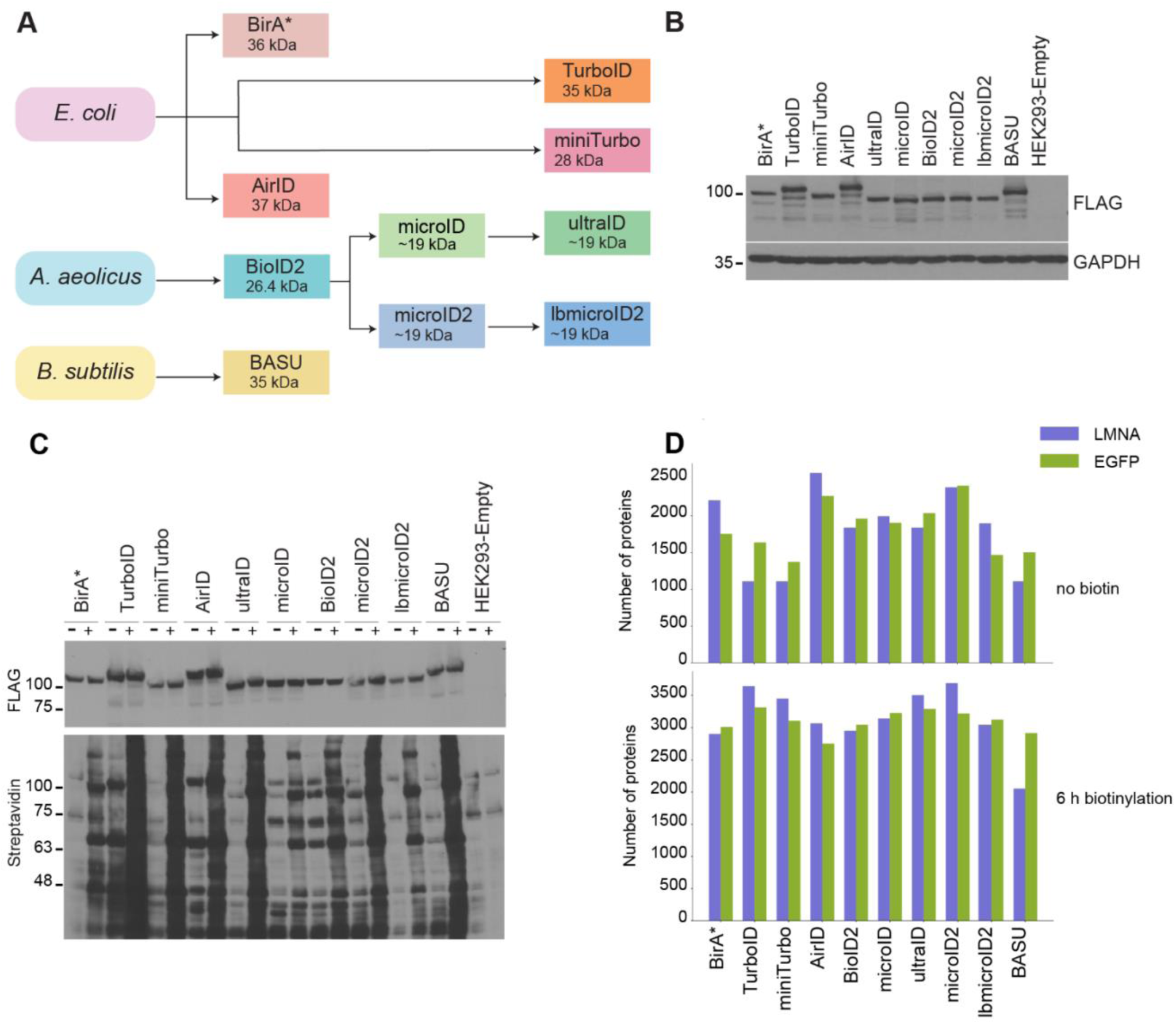
Systematic comparison of ten biotin ligases using LMNA. **A.** Schematic diagram of the BioID enzymes used in this study; enzymes are color-coded based on the bacterial species from which they are derived, and their molecular weights are indicated. **B.** Western blot analysis of HEK293 cells stably expressing LMNA N-terminally fused to different biotin ligases using an anti-FLAG tag antibody. GAPDH is used as a loading control. **C.** Anti-FLAG Western (top) and streptavidin Far-Western (bottom) analysis of HEK293 cells stably expressing LMNA N-terminally fused to various biotin ligases without (-) or with (+) the addition of exogenous biotin for 6 h. **D, E**. Total number of proteins identified by mass spectrometry before SAINTexpress scoring using LMNA- and EGFP-fused biotin ligases after incubation with biotin for 6 h (D) or in the no-added-biotin condition (E) (averaged across replicates).

To compare enzyme efficiency, each biotin ligase was fused N-terminally to the coding sequence of the nuclear lamina component lamin A (LMNA), whose proximal interactors are well characterized^6, 25, 43, 44^. As a negative control, we cloned N-terminally tagged eGFP. Stable pooled transfectants were generated in HEK293 Flp-In T-REx cells and expression was induced by doxycycline (1 µg/mL) for 24 h. Immunoblotting with an anti-FLAG antibody confirmed comparable steady-state expression levels of LMNA across all biotin ligase variants (Fig. 1b), allowing for a comparative analysis.

### Assessment of labeling efficiency at 6 h after biotin addition

To compare the ability of each enzyme to induce biotinylation of proteins proximal to LMNA, we analyzed cells (grown in standard media supplemented with fetal bovine serum) that were either left untreated or incubated for 6 h with 50 µM exogenous biotin—a timepoint and biotin concentration previously established as sufficient for the first-generation BioID enzymes^14, 45^. Far-western blotting using streptavidin-HRP detected biotinylation for all enzymes in the presence of biotin, with TurboID, miniTurbo, ultraID, microID2, and BASU producing the strongest signals (Fig. 1c). In the absence of supplemented biotin, a much lower signal was detected, with the exception of AirID and TurboID, which displayed a clear signal similar to BirA* plus biotin.

Starting with the same amount of lysate, we next performed proximity-dependent biotinylation followed by streptavidin pull-down and mass spectrometry using LMNA and eGFP fusions for each ligase with and without biotin (in triplicate). Across enzymes, ∼1,000–2,500 proteins were identified without biotin supplementation and ∼2,000–3,500 in the biotin-supplemented condition for LMNA or EGFP fusions (Fig. 1d; Supplementary Table 2). To define high-confidence proximal proteins, we applied Significance Analysis of INTeractome (SAINTexpress; Supplementary Table 3; see Methods) with a Bayesian False Discovery Rate (BFDR) cutoff of ≤ 1%^46^, comparing each LMNA fusion to its corresponding eGFP control to account for enzyme-specific background (Supplementary Fig. 1a).

In the presence of biotin, the number of high-confidence proximal interactions ranged from 109 to 338, with TurboID, ultraID, and miniTurbo yielding the most and BirA* and BASU the fewest (Fig. 2a and Supplementary Table 4). In the absence of supplemented biotin, the number of high-confidence interactions varied dramatically across enzymes, ranging from just 3 (lbmicroID2) to over 100 with BioID2 (118), AirID (130), and TurboID (132) (Fig. 2a). Although high sensitivity is often desirable in proximity labeling experiments, the contrast between biotin supplementation and no-added-biotin conditions is a key indicator of labeling inducibility under our experimental conditions. This enrichment was most pronounced for lbmicroID2, followed by miniTurbo, BirA*, and ultraID when comparing the number of high-confidence interactors (Supplementary Fig. 1b). Similar trends were observed when comparing spectral count enrichment for shared high-confidence proteins between conditions (Supplementary Fig. 1c).

**Fig. 2.**
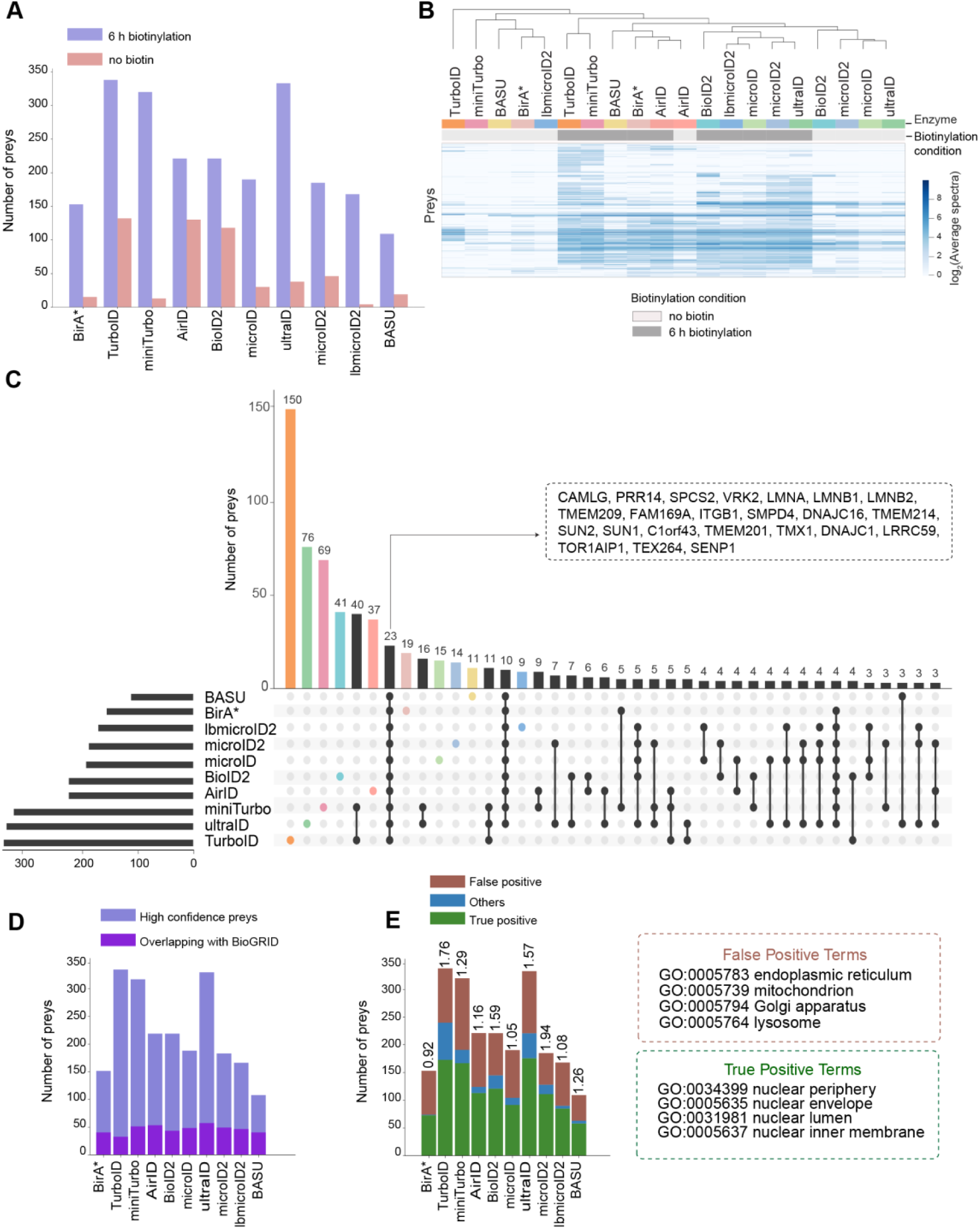
Comparison of prey identification and specificity across biotin ligases using LMNA. **A.** Number of high-confidence preys (SAINTexpress ≤ 1%) identified with or without biotin supplementation (6 h). **B.** Heatmap of log₂-transformed average spectral counts for high-confidence preys across ligases and biotinylation conditions. Samples were clustered using Pearson correlation and average linkage. **C.** UpSet plot showing shared and unique high-confidence preys across ligases. Each row indicates the contributing ligases for a given intersection (black dots). Top bar chart: intersection sizes; side bar chart: total preys per ligase. Preys shared by all ten ligases (n = 23) are listed. **D.** Number of high-confidence preys overlapping with known LMNA interactors from BioGRID. **E.** Subcellular classification of high-confidence preys based on Gene Ontology: Cellular Component annotations. Preys are categorized as true positives (green), false positives (brown), or others (blue) based on predefined localization terms.

Together, these data provide a benchmark for comparing labeling efficiency and inducibility across biotin ligases under standardized conditions, noting that we did not deplete biotin from the serum for these experiments (unlike other studies^42^; see Discussion).

### Comparison of prey profiles across ligases

To assess similarities in labeling patterns, we performed unsupervised hierarchical clustering of high-confidence preys based on their average spectral counts across biological replicates for both biotin and no-added-biotin conditions (Fig. 2b). Enzymes clustered primarily by species origin: all *A. aeolicus*-derived enzymes (ultraID, microID, microID2, lbmicroID2, and BioID2) formed a distinct group, while TurboID and miniTurbo clustered together. For most enzymes, the biotin-supplemented condition was clearly distinct from the standard growth media conditions. A notable exception was AirID, whose biotin and no-added-biotin profiles clustered together—suggesting that substantial background biotinylation in the absence of supplemented biotin may limit its utility for detecting condition-specific interactomes.

We next examined the overlap of high-confidence proximal interactions across enzymes (Fig. 2c). This revealed 23 proteins consistently identified across all ligases, 16 of which localize to nuclear envelope-associated compartments. This core shared interactome included B-type lamins (LMNB1 and LMNB2), which co-assemble with LMNA^47^; LINC complex proteins SUN1 and SUN2, which bridge the nuclear lamina to the cytoskeleton^48, 49^; and nuclear envelope transmembrane proteins TMEM201 and TMEM209. We also detected regulatory factors such as TOR1AIP1 (LAP1), which tether the nuclear lamina to the inner nuclear membrane^44, 50^; VRK2A, whose nuclear envelope retention depends on A-type lamins^51^; and PRR14, which contains a lamina-binding domain mediating its dynamic association with the nuclear lamina^52^. Additionally, we identified FAM169A, a component enriched at the lamina– inner nuclear membrane interface and first discovered using BioID^6^. These comparisons highlight that all enzymes can capture lamina proximal interactors but reveal differences in both prey recovery and background profiles among enzymes.

### Benchmarking against known interactors and GO annotations

To benchmark enzyme performance, we compared high-confidence preys against a curated reference set from BioGRID^53^, filtered for interactions supported by at least two independent experimental lines of evidence (see Methods). All enzymes identified a similar number of known LMNA interactors, with ultraID capturing the highest number (55) (Fig. 2d; Supplementary Table 5). We further assessed prey list quality using a Gene Ontology (GO) term–based classification system. “True” positive terms were selected based on established LMNA localization at the nuclear lamina. Conversely, we defined four cellular compartments spatially separated from the nuclear lamina as “false” positive terms. ultraID, TurboID, and miniTurbo identified the highest number of true positive interactions under these conditions (Fig. 2e; Supplementary Table 6). Among those, specificity (assessed true/false terms) was highest for TurboID (ratio 1.77), followed by ultraID (1.57) and miniTurbo (1.29). These analyses identified ultraID, miniTurbo, and TurboID as the most effective enzymes for mapping the LMNA interactome at 6 h, balancing sensitivity and specificity.

### Assessment of early biotinylation activity across enzymes

We next explored shorter biotinylation times, starting with far-western blots, to identify enzymes appropriate for more time-limited temporal studies. At 5 min, TurboID yielded the strongest signal, followed by ultraID, AirID, and miniTurbo (Supplementary Fig. 1d–f). At 15 min, the same four enzymes also displayed the strongest signals, followed by BASU. At 1 h, the five top enzymes from the 15-min time point still generated the strongest signal, alongside microID.

We selected ultraID, miniTurbo, and TurboID (the enzymes with the highest numbers of proximal interactors at 6 h), as well as BASU (the sole *B. subtilis*–derived ligase), for follow-up mass spectrometry experiments at 5, 15, and 60 min, alongside no-added-biotin controls, and performed SAINTexpress scoring against matched enzyme and timepoint as described above.

Unsupervised hierarchical clustering of the high-confidence proximal interactors across all time points revealed that samples grouped primarily by enzyme (Fig. 3a). The no-added-biotin conditions clustered together for ultraID, miniTurbo, and BASU, consistent with the low signal observed in the absence of biotin. In contrast, the no-added-biotin condition for TurboID clustered with the biotin-treated samples from all enzymes, further supporting its substantial background labeling in standard media even without biotin supplementation.

**Fig. 3.**
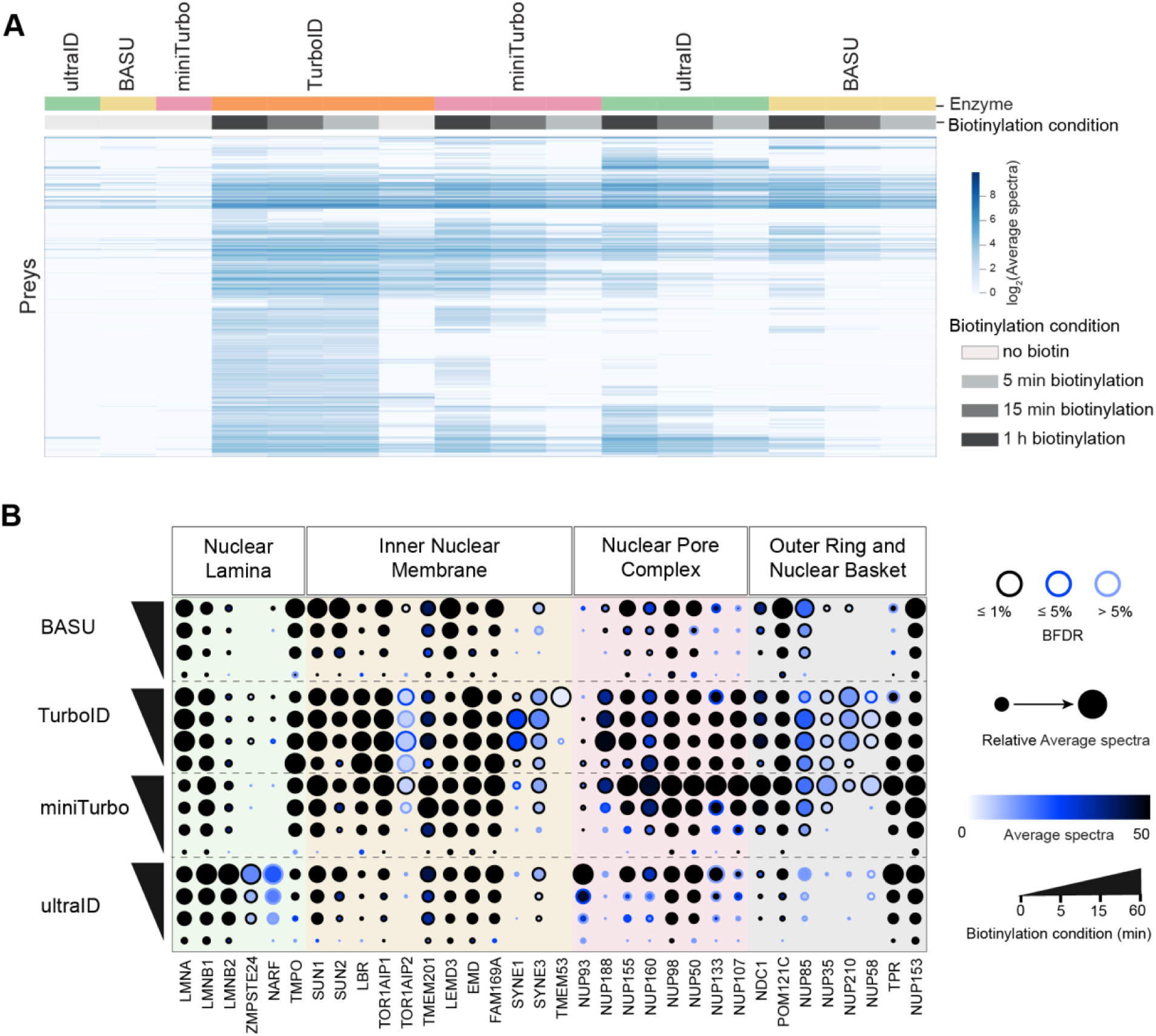
Evaluation of shorter biotinylation times for ultraID, miniTurbo, TurboID and BASU fused to LMNA. **A.** Heatmap of log2-transformed average spectral counts for preys identified with LMNA fused to indicated enzymes and time points (see legend inset). Average spectral counts across replicates were clustered using Pearson correlation and average linkage. **B.** Dot plot showing nuclear lamina, inner nuclear membrane, and nuclear pore complex components (a subset of the proximal interactome) identified with LMNA fused to the different enzymes at the indicated times.

We then examined recovery of nuclear envelope–associated proteins, consistent with LMNA localization (Fig. 3b). All enzymes successfully recovered key nuclear lamina-associated proteins, including LMNB1 and TMPO, many of which were detected at the earliest time points. Inner nuclear membrane preys, such as the LEM-domain proteins LEMD3 and EMD, were identified across all enzymes. Similarly, SYNE3 (Nesprin-3), a KASH domain protein forming the LINC complex with SUN proteins to connect the nuclear lamina to the cytoskeleton^48, 54^, was consistently detected at the 1-h timepoint by all enzymes.

Notable differences in prey recovery were also observed. For example, ultraID and, to a lesser extent, TurboID, captured ZMPSTE24, a known direct LMNA interactor involved in prelamin A maturation^55^. While different nuclear pore subunits were recovered by all enzymes, ultraID yielded the highest spectral counts for NUP93, a structural component that anchors the nuclear pore complex to the nuclear lamina via direct interactions with A-type lamins^21^. The other enzymes, and in particular miniTurbo and TurboID, preferentially recovered outer ring and nuclear basket components.

Beyond the nuclear envelope, we observed that miniTurbo and TurboID preferentially recovered proteins from the ER lumen and ER-Golgi interface—patterns less prominent for ultraID and BASU. To assess spatial specificity more directly, we calculated the ratio of spectral counts for high-confidence “nuclear lumen” proteins versus other compartments (Supplementary Table 7, Methods). Across all time points, while TurboID had the highest total spectral count for nuclear lumen targets, ultraID exhibited the greatest selectivity (e.g., ratio of 2.84 at 15 min), followed by BASU, miniTurbo, and TurboID (e.g., ratio of 1.56 at 15 min).

Together, these results demonstrate that all enzymes are capable of recovering relevant LMNA proximal partners within as little as 5 minutes. While TurboID yielded the highest number of confident preys, its high background labeling in standard media may limit its utility for experiments requiring precise temporal resolution. As such, TurboID was excluded from subsequent experiments.

### Comparing biotin ligases across different cellular compartments

While the results from LMNA experiments indicated that ultraID and miniTurbo (and to a lesser extent BASU) performed well, we next assessed their broad applicability across diverse subcellular compartments. To do so, we expanded our analysis to include baits we previously profiled using BirA* or TurboID: MAPRE3 (microtubule-associated), HIST1H2BG (chromatin-associated), hnRNPA1 (paraspeckle), PDHA1 (mitochondrial matrix)^18^, and EGFR (plasma membrane)^41^. We also included ST6GALNAC1, a Golgi-resident sialyltransferase^56^, to determine whether any of these enzymes could support proximity-dependent biotinylation in the Golgi lumen—a compartment that has historically been challenging to profile with BirA*^18^.

All baits and enzymes were assessed 15 min after the addition of biotin (based on the results with LMNA above), using the same approaches. Note that all baits profiled with the same enzyme were analyzed jointly with SAINTexpress. Unsupervised hierarchical clustering of the high-confidence preys revealed clustering by bait rather than by enzyme, as expected (Fig. 4a). The top GO terms for each cluster matched the expected subcellular localization of their respective bait, with the exception of ST6GALNAC1, which enriched for “ER lumen”. Analysis of high-confidence preys (Fig. 4b) revealed that while all biotin ligases had comparable bait abundance based on spectral counts, their number of high-confidence proximal interactors varied, with ultraID outperforming miniTurbo and BASU in overall prey recovery at that time point.

**Fig. 4.**
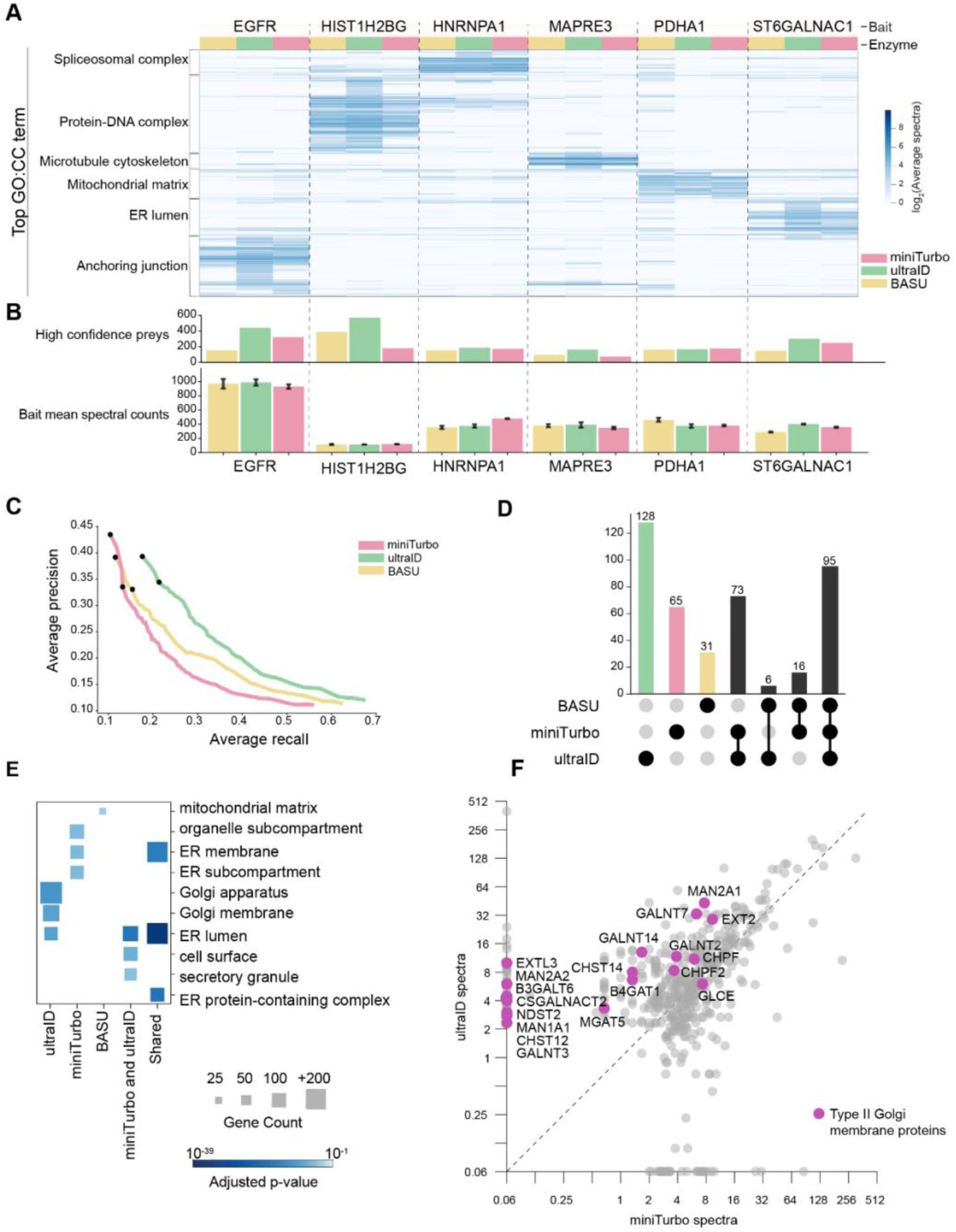
Biotin ligase-specific labeling across subcellular compartments. **A.** Heatmap of log₂-transformed average spectral counts for high-confidence preys across six baits fused to miniTurbo, BASU, or ultraID. Preys were clustered using Pearson correlation and average linkage. Top enriched Gene Ontology: Cellular Component (GO:CC) terms (filtered for categories with ≤1500 genes) annotate major clusters. **B.** Bar plots showing the number of high-confidence preys (top) and average spectral counts of the bait protein (bottom) for each bait-enzyme pair. Error bars are Standard Error of the Mean, SEM (see Methods). **C.** Precision-recall (PR) curves for each ligase across all baits. Ground-truth was defined by relevant GO:CC annotations. Preys were ranked by SAINT BFDR, and curves were generated across BFDR thresholds (0–100% in 1% increments). On each curve, the top dot indicates a BFDR of 1%, and the bottom dot indicates a BFDR of 5%. **D.** UpSet plot showing overlap of high-confidence preys (BFDR ≤ 1%) identified for ST6GALNAC1 with each ligase. **E.** GO:CC enrichment analysis of ST6GALNAC1-associated preys. Top three enriched terms per enzyme or enzyme combination are shown (filtered for categories with ≤1500 genes). Square size reflects gene count; color indicates adjusted p-value. **F.** Scatter plot comparing log₂ average spectral counts for preys identified with ST6GALNAC1 fused to miniTurbo vs. ultraID. Golgi-localized and Type II Golgi membrane proteins are highlighted. Dashed line marks equal abundance.

To comprehensively evaluate enzyme performance beyond prey counts, we performed a precision-recall (PR) analysis using pre-defined reference lists based on GO Cellular Component (CC) terms relevant to each bait’s expected localization (Fig. 4c). Across the dataset, ultraID generally maintained higher precision at equivalent recall points, suggesting that it better recovered true proximity associations while minimizing false positives overall. To examine the specificity of prey recovery, we analyzed high-confidence preys unique to one enzyme, or shared across all, for each bait, followed by GO:CC enrichment analysis. The top 5 GO:CC terms are shown for each group (Supplementary Fig. 2a; Supplementary Table 8). Enriched terms in the shared-across-all-enzymes category recovered biologically relevant annotations for all baits, while the ultraID-only category consistently recovered preys corresponding to meaningful terms. While the results presented above indicate a marked improvement for ultraID over BASU and miniTurbo, when a binning approach was used to compare the prey lists with equal number of ranked preys, the curves from three enzymes were more similar to one another suggesting that they all recover relevant proximity partners overall, albeit in different amounts (Supplementary Fig. 2b). Notably, at a recall of 0.5—where the true signal appears to plateau—the areas under the curve were comparable across enzymes, with ultraID at 0.110, BASU at 0.100, and miniTurbo at 0.085.

The Golgi lumen (exemplified here by the ST6GALNAC1 bait) could not be mapped successfully in our earlier large-scale proximal biotinylation study^18^. Although the shared ST6GALNAC1 cluster was enriched for “ER lumen” (Fig. 4a), manual inspection revealed a substantial number of Golgi lumen proteins across the dataset. We categorized high-confidence preys as shared across all enzymes, unique to one enzyme, or shared between pairs of enzymes (Fig. 4d). GO enrichment analysis was performed for each of these lists, with the 3 top GO:CC (Cellular Component) terms highlighted. While the shared preys (and those unique to miniTurbo) predominantly recovered ER lumen terms (consistent with Golgi enzyme biosynthesis and trafficking), ultraID-unique preys were instead enriched for “Golgi apparatus” and “Golgi membrane” terms, suggesting improved localization and labeling performance by this enzyme (Fig. 4e). Note that BASU-only enriched for a single term, mitochondrial matrix.

We next performed a pairwise comparison of spectral counts for miniTurbo and ultraID fusions across all high-confidence prey proteins, highlighting Type II Golgi membrane proteins, which share the same topology as ST6GALNAC1 (Fig. 4f; Supplementary Fig. 2c). Consistent with the GO enrichment, ultraID consistently achieved higher spectral counts for multiple glycosyltransferase families, including GalNAc-transferases (GALNT2, GALNT3, GALNT7, GALNT14), which initiate O-glycosylation, as well as mannosidases (MAN1A1, MAN2A1, MAN2A2), glycosaminoglycan enzymes (CHPF, CHPF2, CHST12, CHST14), and other glycan-modifying enzymes (B3GALT6, B4GAT1, CSGALNACT2). These results confirm that ultraID is capable of labeling Golgi-resident proteins, a compartment that had proved challenging in previous investigations^18^.

### Comparison of miniTurbo and ultraID across cytoskeletal, endosomal, and cell junctional baits in HeLa cells

To validate and extend our findings in a different cellular context, we next compared ultraID and miniTurbo in HeLa cells using eight proteins spanning major structural elements—actin filaments (ACTB, PFN1, VASP), intermediate filaments (KRT8), endosomal compartments (RAB5A, RAB11A), and plasma membrane/junctional components (RHOB, OCLN)—with the rationale that cytoskeletal components may be particularly sensitive to fusion-induced misfolding or mislocalization^57^.

We first assessed bait expression levels by western blot analysis to ensure a fair comparison between the miniTurbo and ultraID fusions. For RAB5A and RAB11A, the ultraID fusions initially showed higher expression, but tetracycline titration enabled similar expression prior to proximity-dependent biotinylation (Supplementary Fig. 3a). Hierarchical clustering of high-confidence preys grouped samples by bait, with further sub-clustering by functional category (Fig. 5a). The endosomal baits RAB5A and RAB11A clustered together and enriched for intracellular transport proteins. The OCLN and RHOB cluster was enriched for anchoring junction components. ACTB, PFN1, and VASP grouped for their enrichment of actin cytoskeleton preys, while KRT8 formed a distinct cluster enriched for intermediate filament proteins. Across most baits, ultraID recovered a higher number of high-confidence preys than miniTurbo (Fig. 5b).

**Fig. 5.**
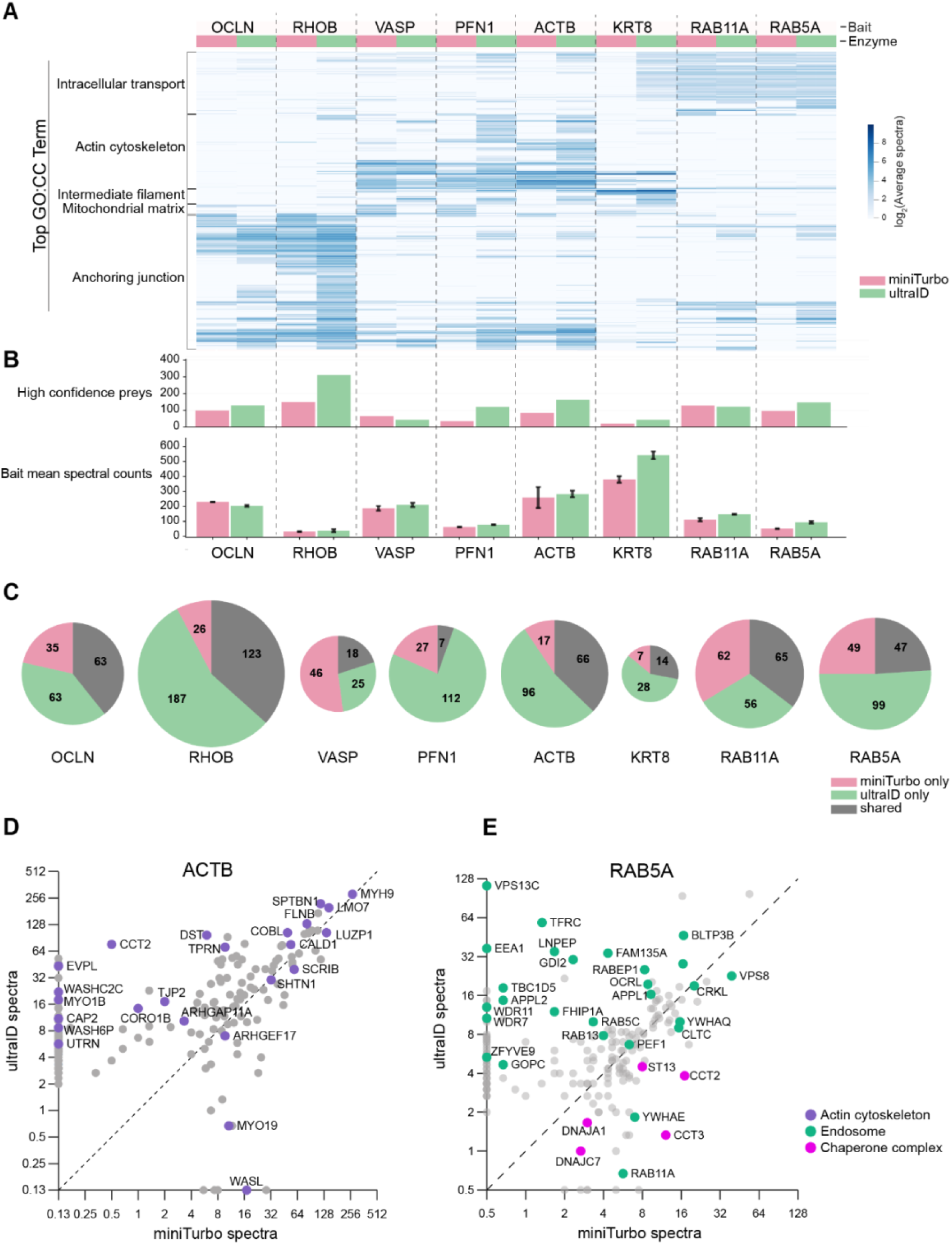
Comparative performance of miniTurbo and ultraID for compartment-specific labeling in HeLa cells. **A.** Heatmap of log₂-transformed average spectral counts for high-confidence preys detected with eight cellular baits fused to miniTurbo or ultraID. Preys were clustered by Pearson correlation and average linkage. Top Gene Ontology: Cellular Component (GO:CC) terms (filtered to categories with ≤1500 genes) annotate major clusters. **B.** Bar plots showing the number of high-confidence preys (top) and average spectral counts of the bait proteins (bottom) for each bait-enzyme pair. Error bars are Standard Error of the Mean, SEM (see Methods). **C.** Pie charts showing the distribution of high-confidence preys (BFDR ≤ 1%) for each bait. Preys uniquely identified by miniTurbo or ultraID are shown in pink and green, respectively; shared preys are shown in gray. **D, E.** Scatter plots comparing log₂-transformed average spectral counts of high-confidence preys identified with ACTB (D) or RAB5A (E) fused to miniTurbo (x-axis) or ultraID (y-axis). The diagonal dashed line indicates equal abundance. Preys are color-coded based on GO:CC annotation.

To examine the specificity of prey recovery, we analyzed high-confidence preys unique to one enzyme, or shared between both, for each bait (Fig. 5c), followed by GO:CC enrichment analysis. The top 3 GO:CC terms are shown for each group (Supplementary Fig. 3b; Supplementary Table 9). For some baits, such as ACTB or RAB11A, there was strong overlap between preys recovered by both enzymes, with concordant GO term enrichment. In other cases—notably for VASP and PFN1—the prey sets were more distinct. For example, ultraID recovered biologically relevant terms (actin cytoskeleton and anchoring junctions) while miniTurbo enriched for unexpected compartments such as the mitochondrial matrix (Supplementary Fig. 3a).

To illustrate these differences, we performed pairwise comparisons of spectral counts (Fig. 5d,e; 6a,b; Supplementary Fig. 4a–d). For ACTB, both enzymes recovered actin cytoskeleton-associated proteins. However, ultraID, which recovered markedly higher numbers of preys, uniquely identified several established actin regulators and junctional proteins, including CAP2, a recycler of actin monomers^58^; CTNNA1, a junctional actin anchor^59^; UTRN, an actin-binding protein^60^; AFDN, a junction-associated scaffolding protein^61^; and CALD1, an actin-associated motor protein^62^, along with many other known actin interactors that were not captured by the miniTurbo variant (Fig. 5d).

**Fig. 6.**
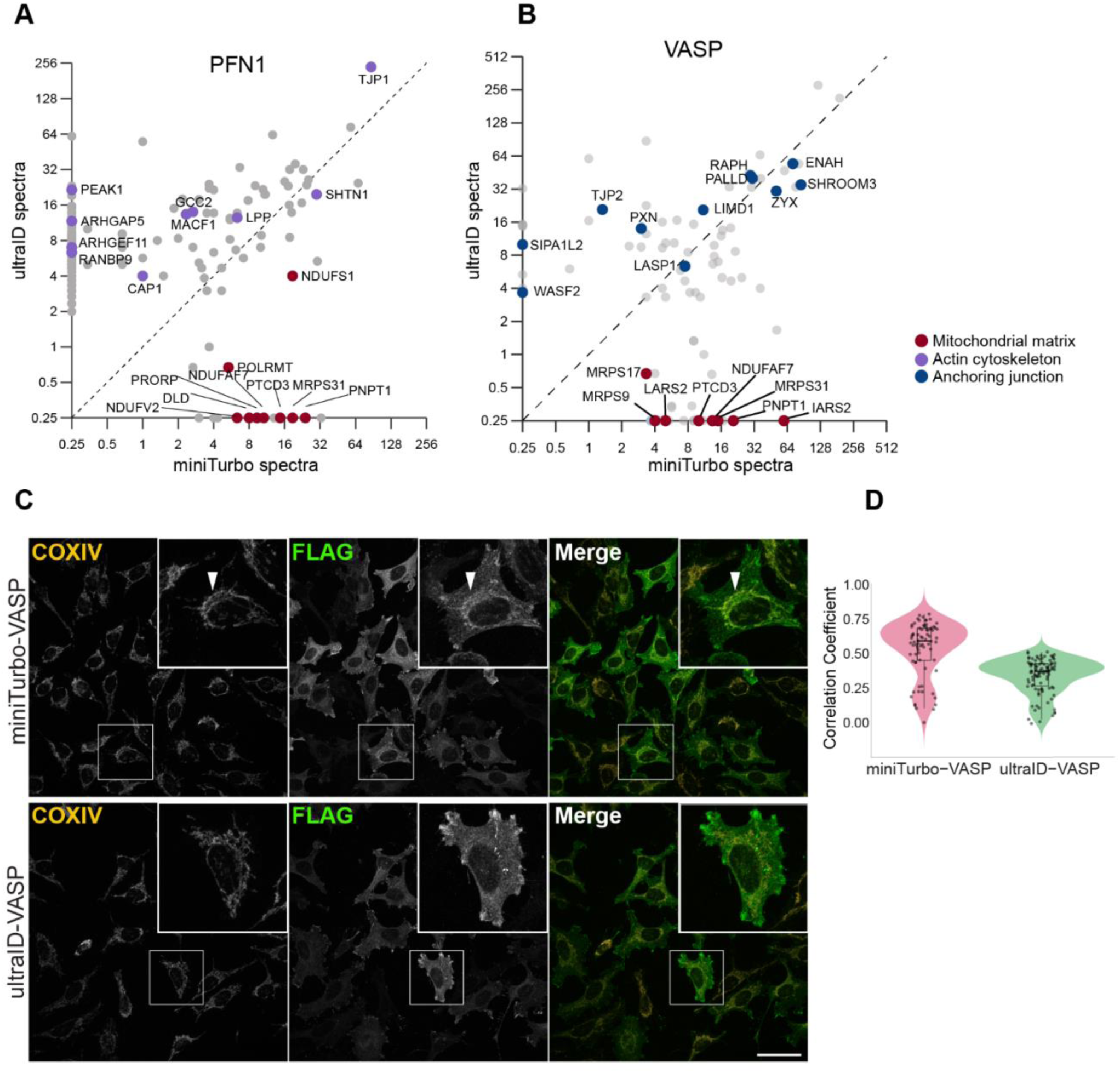
Potential mislocalization of miniTurbo fusions to mitochondria. **A, B.** Scatter plots comparing the abundance of high-confidence proximity interactors for PFN1 (A) and VASP (B) fused to miniTurbo (x-axis) or ultraID (y-axis), based on log₂-transformed average spectral counts across n = 3 replicates. The diagonal dashed line indicates equal abundance between the two fusions. Preys are color-coded by GO:CC annotation. **C.** Colocalization analysis of VASP fusions with mitochondria. HeLa cells stably expressing miniTurbo-VASP and ultraID-VASP were fixed and probed with an antibody against COXIV (mitochondrial marker, yellow) and exogenous VASP using anti-FLAG antibody (green). White arrowheads indicate a representative example of colocalized foci. Scale bar = 25 μm. **D**. Quantification of colocalization between FLAG-tagged VASP fusions and the mitochondrial marker COXIV using Pearson’s correlation coefficient. Violin plots show the distribution of per-cell correlation coefficients (each point represents one cell; n = 76 for miniTurbo-VASP, n = 98 for ultraID-VASP). See Supplementary Table 10.

RAB5A, a small GTPase critical for early endosome fusion and trafficking^63, 64^, recovered endosomal-related proteins, including BLTP3B, VPS8, APPL1, RAB13, YWHAQ, CLTC, and OCRL, when fused with either enzyme, though many of them were quantitatively enriched with ultraID (Fig. 5e). Furthermore, ultraID uniquely identified several known RAB5A interactors that were not detected by miniTurbo, including d EEA1^65^, APPL2^66^, and ZFYVE9. Conversely, miniTurbo-RAB5A showed higher spectral counts for chaperonin complex proteins, such as CCT2, ST13, CCT3, DNAJA1, and DNAJC7, compared to ultraID-RAB5A, potentially indicating folding issues with the miniTurbo-RAB5A fusion (note that chaperonin recovery was also more prominent with miniTurbo-RAB11A than ultraID-RAB11A, Supplementary Fig. 4a). When fused to RHOB, despite the recovery of nearly twice as many confident preys with ultraID, both enzymes identified preys related to anchoring junctions and the plasma membrane, suggesting a similar specificity (Supplementary Fig. 4b). Occludin (OCLN) showed more similar number of recovered preys and similar GO term enrichment, but some quantitative differences in the preys recovered (Supplementary Fig. 4c). Lastly, ultraID yielded a higher number of high-confidence biologically relevant preys (Supplementary Fig. 4d).

### Aberrant localization to the mitochondria is associated with some of the miniTurbo fusions

While analyzing PFN1 and VASP, key regulators of actin dynamics^67, 68^, we observed that miniTurbo-based proximity labeling unexpectedly recovered a large number of mitochondrial proteins, inconsistent with their known cytoskeletal localization (Fig. 6a,b). Scatter plot visualizations highlight this differential recovery: miniTurbo-PFN1 and miniTurbo-VASP uniquely identified multiple mitochondrial proteins, including IARS2, GRSF1, PNPT1, NDUFS1, FASTKD2, MRPS31, PTCD3, and HSPE1, compared to ultraID fusions. By contrast, ultraID-PFN1 and ultraID-VASP recovered preys enriched in anchoring junction and actin cytoskeleton components, aligning with the established localizations of these proteins.

These results suggested potential mislocalization of the miniTurbo fusions. To determine whether the observed mitochondrial enrichment was due to mislocalization, we conducted immunofluorescence microscopy using a mitochondrial marker (COXIV). In contrast to ultraD-VASP, which exhibited the expected cytoplasmic and plasma membrane-associated distribution, miniTurbo-VASP showed partial colocalization with mitochondria in a subset of cells (Fig. 6c). Quantitative analysis of approximately 100 cells revealed that 73% of miniTurbo-VASP-expressing cells exhibited a Pearson correlation coefficient of at least 0.4 with the mitochondrial marker, compared to only 8% of ultraID-VASP-expressing cells (Fig. 6d; Supplementary Table 10), supporting a significant enrichment of miniTurbo-VASP at mitochondria.

These findings support the hypothesis that aberrant localization of miniTurbo fusion proteins may contribute to the nonspecific enrichment of mitochondrial proteins and impact the interpretation of proximity labeling data in some contexts.

## Discussion

Our systematic investigation of biotin ligase variants revealed that all enzymes tested recovered relevant partners for LMNA, consistent with the fact that this bait was used for the development of BioID and the benchmarking of several of the other enzymes. For most baits tested across two or more enzymes, the profiles aligned with the reported subcellular localization of the bait, supporting the view that proximity labeling with any of these enzymes can yield useful biological insights, as has been widely reported in the literature. That said, our results also show that enzyme choice can influence the results, especially in cases where bait behavior, localization, or fusion stability varies across constructs.

As previously reported ^29, 42^, TurboID is highly active but supports substantial labeling even in the absence of supplemented biotin, limiting its utility for time-resolved experiments. While we used standard Fetal Bovine Serum (FBS), prior work has shown that TurboID continues to label even after several days in dialyzed FBS^42^, likely due to intracellular biotin pools. AirID exhibited a similar pattern, with profiles in the presence and absence of biotin clustering together, indicating limited dynamic range. The other enzymes were more responsive to exogenous biotin, but BirA*, BioID2, microID, microID2, and lbmicroID2 did not achieve the same level of labeling efficiency or speed as TurboID, AirID, miniTurbo, ultraID, and, to a lesser extent, BASU.

Among the enzymes tested, ultraID consistently performed well across subcellular compartments, labeling durations, and expression contexts. It efficiently captured known interactors of nuclear lamina, cytoskeletal, endosomal, and Golgi-resident proteins. In direct comparisons with miniTurbo and BASU, ultraID demonstrated both higher prey recovery and greater spatial specificity, particularly for the Golgi-resident bait ST6GALNAC1. Similarly, ultraID outperformed miniTurbo in recovering actin- and junction-associated proteins, using PFN1 and VASP as baits, whereas miniTurbo enriched for mitochondrial proteins, consistent with partial mislocalization confirmed by immunofluorescence. The partial mislocalization to mitochondria in miniTurbo fusions was also observed for some baits in our study of the SARS-CoV-2 proximal interactome^69^, suggesting that this phenomenon may be more common than previously appreciated. In most of the comparisons, both miniTurbo and ultraID outperformed BASU in terms of relevant prey recovery.

Our study has several limitations. We benchmarked fewer than 20 baits and did not examine all cellular compartments or membrane topologies. Broader proteome-scale comparisons in additional cell types or organisms will be important to fully define enzyme-specific performance features. While we elected to perform comparisons at the same time points across enzymes, this may have negatively affected the results with the first-generation enzymes (e.g., BirA* and BioID2, in the 6 h labeling experiments), as well as some of the later comparisons between TurboID, miniTurbo, ultraID, and BASU at later timepoints. Furthermore, each enzyme exhibits a distinct background labeling profile, necessitating the use of matched controls for accurate prey scoring. It is also likely that preys that are labeled non-specifically by a specific enzyme would be challenging to identify as true proximal partners with that enzyme; in these cases, selecting a different enzyme may be helpful. Although a comprehensive background profiling effort was beyond our scope, the patterns observed in this dataset—together with the high-confidence preys from eGFP controls provided in Supplementary Table 2 offer a resource for enzyme selection and experimental design in proximity proteomics.

In conclusion, our benchmarking study demonstrates that proximity-dependent biotinylation enzymes differ substantially in labeling efficiency, inducibility, and spatial specificity. While all tested enzymes can identify proximal interactors under appropriate conditions, their performance varies by labeling duration, subcellular context, and fusion behavior. Among the variants examined, ultraID offered the most consistent combination of low background, rapid labeling, and robust prey recovery across the baits and compartments tested. These features make it a strong candidate for applications requiring short labeling times or higher spatial fidelity. However, other enzymes—including TurboID and miniTurbo—may remain preferable in contexts prioritizing maximal labeling efficiency or in organisms where ultraID performance has not yet been validated. The accompanying datasets and controls provide a practical resource to inform enzyme selection and guide experimental design in proximity proteomics.

## Acknowledgements

We thank all members of the Gingras laboratory, particularly Cassandra J. Wong and Drs. Rasha Al Mismar, Ugo Dionne, and Jonathan Roth for their advice on the project. We are grateful to Drs. Brian Raught and Geoffrey Hesketh for critical review of the manuscript, and Dr. Laurence Pelletier for advice on the microscopy. This work was funded by grants from the Canadian Institutes of Health Research (CIHR PJT-185987), the Terry Fox Research Institute (TFRI; PPG 1107), and the Natural Sciences and Engineering Research Council of Canada (NSERC RGPIN-2019-06297) to A.-C.G. A.-C.G. is the Canada Research Chair (Tier 1) in Functional Proteomics and the Dr. Louis Siminovitch Chair in Research. J.K. was supported by a Canada Graduate Scholarship—Doctoral from the Natural Sciences and Engineering Research Council of Canada. S.S and K.V were supported by University of Toronto Open Fellowships. The mass spectrometry data were acquired at the Proteomics Facility (RRID: SCR_025375), and the microscopy was performed at the High-Content Screening Facility (RRID: SCR_025391) at the Network Biology Collaborative Centre, Lunenfeld-Tanenbaum Research Institute. The Centre is supported by the Canada Foundation for Innovation and the Government of Ontario.

## Author Contributions

A-C.G., W.R.H., S.S., and K.V. conceptualized the study. W.R.H. and R.K. designed the cloning vectors. W.R.H., R.K., Q.H., S.S., and K.V. performed cloning. S.S. performed HEK293-based transfections, selection, and harvesting, with Z.Y.L. performing the pull-down assays. K.V. was responsible for all HeLa-based experiments. S.S., K.V., V.K., J.K. performed data analysis. S.S., K.V., and A-C.G. interpreted the biological relevance. M.H. provided expertise in microscopy image analysis. B.S. provided expertise in mass spectrometry data analysis. S.S., K.V., and A-C.G. wrote the manuscript with input from all authors. A-C. G supervised the project.

## Online methods

### Plasmids

Destination (DEST) plasmids in the pcDNA5/FRT/TO backbone (Thermo Fisher Scientific, Invitrogen, V652020) for expression of BirA*^6^, BioID2^25^, miniTurbo^29^ and TurboID^70^ fusions were reported previously. Open Reading Frames (ORFs) encoding ultraID^26^, AirID^28^, BASU^27^, microID^26^, microID2 and lbmicroID2^24^ were codon-optimized for expression in human cells and synthesized by Twist Bioscience (see full sequences in Supplementary Table 1). Two Gateway-compatible pcDNA5/FRT/TO-DEST plasmids were generated for each variant to enable either N-terminal or C-terminal bait tagging. pcDNA5/FRT/TO-DEST plasmids for N-terminal tagging were constructed by excising the miniTurbo ORF from pcDNA5/FRT/TO-N-term-miniTurbo-DEST as a KpnI/NsiI restriction fragment and replacing it with a KpnI/NsiI-digested restriction fragment containing the variant ORF. pcDNA5/FRT/TO-DEST plasmids for C-terminal tagging were created by excising the 3xFLAG-miniTurbo ORF from pcDNA5/FRT/TO-C-term-miniTurbo-DEST as a KpnI/XhoI restriction fragment and replacing it with a KpnI/XhoI-digested restriction fragment containing the 3xFLAG-variant ORF. Bait ORFs were introduced into pcDNA5/FRT/TO-DEST-enzyme via Gateway cloning following the manufacturer’s instructions (Thermo Fisher Scientific). All constructs were validated by restriction digest analysis and Sanger sequencing of the subcloned fragments.

### Cell culture, stable cell line generation and bait expression monitoring

HEK293 Flp-In T-REx (Invitrogen, Cat # R78007) cells were maintained in DMEM supplemented with 10% fetal bovine serum (FBS), 100 U/mL penicillin, and 100 μg/mL streptomycin. Flp-In T-REx HeLa cells (a kind gift from Dr. Arshad Desai, Ludwig Cancer Research, La Jolla, USA) were cultured in DMEM containing 5% FBS, 5% Cosmic Calf Serum (CCS), 100 U/mL penicillin, and 100 μg/mL streptomycin. Stable cell lines were generated as previously described^71^ and maintained in media supplemented with 200 μg/mL hygromycin B. Protein expression was induced in HEK293 Flp-In T-REx by adding 1 µg/mL doxycycline (Dox) and in HeLa cells by adding 1 µg/mL tetracycline for 24 h (unless otherwise described). Cells were routinely tested for mycoplasma contamination (MycoAlert kit, Lonza, Cat# LT07), but were not independently authenticated.

Cells were seeded at a density of 1 × 10⁶ cells/mL in 10 cm plates. Each bait condition was performed in triplicate. Protein expression was induced 24 h after seeding, and biotinylation was initiated 24 h later (at ∼90% confluence) by adding 50 μM biotin for the indicated durations. Cells were lysed in SDS-based buffer (25 mM Tris-HCl, pH 7.4, 2% SDS), heated in Laemmli sample buffer, and subjected to SDS-PAGE. Proteins were transferred onto nitrocellulose membranes (GE Healthcare, Cat# 10600001) for immunoblot analysis. After Ponceau S staining, membranes were blocked in 3% non-fat milk in TBST and probed with mouse anti-FLAG antibody (Sigma-Aldrich, Cat# F3165, 1:2000), followed by anti-mouse IgG-HRP (GE Healthcare, Cat# NA931, 1:5000). Streptavidin staining was performed using HRP-conjugated streptavidin (GE Healthcare, Cat# RPN1231vs, 1:2500) after blocking in 3% BSA. Rabbit mAb anti-GAPDH (14C10, Cell Signaling Technology, Cat# 2118, 1:2000) served as the loading control, followed by HRP-conjugated rabbit IgG (Cytiva, Cat# NA934, 1:10,000). Membranes were visualized using LumiGLO (Cell Signaling Technology, Cat# 7003S) or ECL reagent (Global Life Sciences Solutions, Cat# RPN2232) and imaged on a BioRad ChemiDoc system.

### BioID sample preparation

Cells were seeded in 10 cm plates and induced with 1 µg/mL doxycycline or tetracycline at approximately 50% confluence for 24 h. Biotinylation was then initiated by adding 50 µM biotin at specific time points (5 min, 15 min, 1 h, and 6 h). Following induction and labeling, cells were washed with cold PBS, scraped into 1 mL PBS using a spatula, and transferred to pre-weighed tubes. After centrifugation at 500 × g for 5 min, the supernatant was removed, and the cell pellets were flash-frozen on dry ice.

After determining the pellet weights, pellets were thawed on ice and lysed with modified RIPA buffer (50 mM Tris-HCl, pH 7.4, 150 mM NaCl, 1 mM EGTA, 0.5 mM EDTA, 1 mM MgCl₂, 1% NP-40, 0.1% SDS, 0.4% sodium deoxycholate, 1 mM PMSF [Bioshop, Cat# PMS123.5], and 1× protease inhibitor cocktail [Sigma-Aldrich, Cat# P8340]) at a 10:1 ratio volume:weight, e.g., 1000 μL buffer for a 1000 μg pellet. Lysates were then sonicated (three cycles: 5 s on, 3 s off, 30% amplitude, Q500 Sonicator, Cat# 4422) and treated with TurboNuclease (1 µL/sample, BioVision, Cat# 9207-50KU) and RNase (1 µL/sample, Sigma) for 15 min at 4 °C. Additional SDS was added to bring the final concentration to 0.4%, followed by another 15-min incubation at 4 °C. Lysates were centrifuged (15,000 × g, 15 min), and supernatants were collected.

Biotinylated proteins were enriched using streptavidin-Sepharose resin (GE Healthcare, Cat# 17511301). Pre-washed beads (20 µL bed volume per sample) were incubated with clarified lysates for 3 h at 4 °C with end-over-end rotation. Following incubation, the beads were washed sequentially: once with SDS wash buffer (25 mM Tris-HCl, pH 7.4, 2% SDS), twice with RIPA buffer, once with TNEN buffer (25 mM Tris-HCl, pH 7.4, 150 mM NaCl, 1 mM EDTA, 0.1% NP-40), and three times with 50 mM ammonium bicarbonate buffer (pH 8.0). For on-bead digestion, beads were incubated with 1 µg trypsin (Sigma-Aldrich, Cat# T6567) in 70 µL ammonium bicarbonate buffer overnight at 37 °C with rotation. A second digestion was performed with 0.5 µg trypsin for 3 h. Peptide supernatants were collected, pooled with two bead washes with HPLC water, centrifuged (10,000 rpm, 2 min), and acidified with 5% formic acid before vacuum drying. Samples were stored at −80 °C and resuspended in 2.5% formic acid for mass spectrometry analysis.

### Immunofluorescence and image analysis

Flp-In T-REx HeLa cells were seeded on 12 mm coverslips (Micro Cover Glass, Electron Microscopy Sciences, Cat# 72230-01) placed in 24-well plates and treated with 1 µg/mL tetracycline for 24 h to induce protein expression. Cells were washed with PBS supplemented with 200 mM calcium chloride and 100 mM magnesium chloride (PBS+/+), then fixed in 4% paraformaldehyde for 15 min at room temperature. Permeabilization was carried out using 0.1% NP-40 in PBS for 10 min, followed by two PBS washes and blocking in 2% nonfat milk in PBS for 60 min at room temperature.

Cells were incubated with anti-FLAG M2 monoclonal antibody (Sigma-Aldrich, Cat# F3165, 1:2000 in 2% nonfat milk) and anti-COX IV (3E11) rabbit monoclonal antibody (Cell Signaling Technology, Cat# 4850S, 1:500 in 2% nonfat milk) for 3 h on ice with gentle rotation. After three PBS washes, cells were incubated with Alexa Fluor 488–conjugated goat anti-mouse secondary antibody (Molecular Probes, Cat# A11005, 1:1000), Alexa Fluor 594–conjugated goat anti-rabbit secondary antibody (Molecular Probes, Cat# A11012, 1:1000), and Hoechst 33342 (Life Technologies, Cat# H3570, 10 mg/mL stock, 1:10000) prepared in 2% BSA in PBS. Incubation was performed at room temperature for 1 h with gentle rotation. Coverslips were washed three times with PBS, mounted on Superfrost Plus glass slides (Fisherbrand, Cat# 1255015) using ProLong Gold Antifade Mountant (Molecular Probes, Cat# P36930), and imaged using a Nikon A1 confocal microscope equipped with a 60× oil immersion objective (NA 1.4).

Images were acquired in DAPI (Ex. 405 nm, Em. 450 nm), FITC (Ex. 488 nm, Em. 525 nm) and TRITC (Ex. 461 nm, Em 595 nm) channels as 18-step z-stacks at 0.5 µm intervals and processed using maximum intensity projection in Nikon NIS-Elements AR software (Version 5.30.01). Acquisition parameters were kept constant across all samples within each experiment to ensure reproducibility.

### Image quantification

Quantification of cytoplasmic fluorescence in the red and green channels was performed using a custom pipeline developed in CellProfiler 4.2.4^72^. After background thresholding, nuclei were segmented in the DAPI channel, and nuclei touching image borders were excluded. Cytoplasmic regions were defined as secondary objects surrounding the primary nuclear objects, identified using the red channel. Colocalization between green and red cytoplasmic signals was then calculated within the red-defined cytoplasmic masks on a per-cell basis and reported as a correlation coefficient (Supplementary Table 10).

### Mass spectrometric analysis

Samples were analyzed using a timsTOF Pro 2 quadrupole time of flight (qToF) trapped ion mobility mass spectrometer (Bruker, Billerica, MA, USA; timsControl 5.0.9, Compass HyStar 6.2.1.13). The instrument was coupled to an Evosep One liquid chromatography (LC) system (Evosep, Odense, Denmark; EvoSepONE driver version 2.2) configured with the 60SPD 22-min standard gradient and Evosep EV1109 Performance columns (8 cm × 150 μm, 1.5-μm C18 packing). For each sample, one-eighth of the tryptic peptide digest was prepared using EvoTip Pure sample tips following the manufacturer’s protocol. The mass spectrometer was operated in a DDA parallel accumulation-serial fragmentation (PASEF)^73^ mode with 10 PASEF frames per cycle, and active dynamic exclusion was selected using a polygonal filter that excluded singly charged species based on their ion mobility. Ion mobility was ramped from 0.6 to 1.6 1/K0 in 100 ms with a matching 100-ms accumulation time, resulting in a total cycle time of 1.16 s. Collision energy for the selected precursors was nonlinearly dependent on ion mobility within the range of 17.12 eV at inverse mobility 0.6 to 76.46 eV at inverse mobility 1.6.

### MS data analysis

Data in Bruker’s “.d” format were searched using FragPipe (version 17^74^) integrated within the ProHits platform^75^ and analyzed with the MSFragger search engine (version 4.1). Searches were conducted against the human UniProt database (UP000005640, updated 26 October, 2021), which was supplemented with common contaminants and tag sequences, including BirA*, TurboID, miniTurbo, ultraID, microID, microID2, lbMicroID2, BioID2, BASU, AirID, and EGFP. The total number of entries was 40,840 including reverse (decoys) sequences. The search was set to identify tryptic peptides, allowing up to two missed cleavages. The peptide and fragment ion mass tolerances were both set to ±40 ppm. Peptide variable modifications were set as acetylation at the protein N-terminus and oxidation of methionine (M).

The peptide-spectrum match (PSM) results from MSFragger were validated through a computational post-processing tool, Percolator, and a deep learning prediction model, MSBooster, with default parameters, which are incorporated through FragPipe^76^. All proteins with 1% FDR were filtered (by default 1% FDR at the PSM, ion, peptide, and protein levels) using Philosopher^77^. Only proteins with protein probability ≥0.95 and containing at least two unique peptides were considered for further analysis.

### Proximal interaction scoring

SAINTexpress^46^ (Version 3.6.3) was used with default parameters to analyze the proteomic data; no bait compression was used (i.e., each of the triplicate experiments was analyzed separately before averaging the results in a single SAINT score used for Bayesian FDR modelling). Negative controls included HEK293 Flp-In T-REx and HeLa Flp-In T-REx cells expressing eGFP fused to the respective biotin ligase, as well as the corresponding parental cell lines without any tagged construct. Analyses were performed separately for each enzyme, time point, and cell line, with each compared against six control runs (three eGFP and three parental). The rationale for doing this analysis separately is that the background contamination profiles (as defined in identifications across the controls) vary by both time point, enzyme and cell line. A single SAINTexpress run for all baits analyzed across a specific set of conditions was performed (i.e., all samples for Figure 4 were jointly analyzed; all samples for Figures 5 and 6 were jointly analyzed). For HEK293 samples, the six individual control runs were used individually for SAINTexpress scoring. For HeLa samples, the six control runs were compressed into three “virtual” controls to maximize stringency in scoring^78^.

Sample reproducibility was assessed through pairwise replicate comparisons after SAINTexpress analysis, and the resulting R² values were reported (Supplementary Table 3).

For bait recovery assessment, error bars represent the standard error of the mean (SEM) calculated from three biological replicates. For each bait, spectral count values from three independent experiments were extracted from the SAINT file, and SEM was calculated as σ/√3, where σ is the standard deviation of the three replicates. For H2BC8, the error was calculated for H2BC12 (a nearly identical protein to which the peptides were assigned in the protein identification step above).

### Evaluation of enzyme performance with BioGRID and Gene Enrichment Analysis

To evaluate enzyme performance, we assessed the overlap between high-confidence preys identified in our dataset and a curated reference set of LMNA interactors obtained from BioGRID v4.4 (Supplementary Table 5). The BioGRID^53^ reference set was filtered to include only interactions supported by at least two independent experimental sources. Overlap analysis results are presented in Fig. 2d as a stacked bar chart.

Gene Ontology (GO) enrichment analysis was performed using g:Profiler^79^, with default parameters and the organism set to Homo sapiens. To assess the biological relevance of BioID-identified preys, we generated a benchmarking dataset using Gene Ontology: Cellular Component (GO:CC) annotations. A comprehensive, non-redundant list of all SAINT-passed preys was compiled and analyzed using g:Profiler to identify associated GO:CC terms. For specific baits—such as LMNA—a list of biologically relevant GO terms (e.g., *nucleus, nuclear periphery*) was curated to define likely true interactors. In parallel, a set of unrelated GO terms (e.g., *mitochondrion*, *Golgi apparatus*) was assembled to flag potential false positives (Supplementary Table 6). Using these benchmarking lists, we developed a classification workflow to evaluate prey identification at a stringent SAINT BFDR threshold (≤ 1%). Preys were categorized as true positives (TP), false positives (FP), or “others.” TP and FP classifications were based on overlap with the benchmarking GO:CC terms. Preys falling outside both lists were grouped into the “others” category, which was further analyzed via GO:CC enrichment in g:Profiler to uncover potentially novel or under-annotated localizations (Supplementary Table 6).

In order to score the specificity of each enzyme in recovering inner nuclear membrane preys, GO:CC term *nuclear lumen* (GO:0031981) was used. Average spectral counts of preys that were in this category were divided by the average spectral count of the rest of the preys and were reported as ratios (Supplementary Table 7).

### Precision-recall analysis

To evaluate the performance of proximity labeling enzymes (ultraID, miniTurbo, and BASU), we generated precision-recall (PR) curves for each bait. Benchmarking lists, microtubule (GO:0005874), Golgi apparatus (GO:0005794), mitochondrial matrix (GO:0005759), chromatin (GO:0000785), plasma membrane (GO:0005886), cell junction (GO:0030054), spliceosomal complex (GO:0005681), and ribonucleoprotein complex (GO:1990904) were defined as proteins sharing GO:CC terms with the bait of interest. In the first approach (Fig. 4c), proximal interactors were ranked by SAINT-derived BFDR scores, and precision and recall were calculated over a full range of BFDR thresholds^46^, from 0 to 100% (in 1% increments). Precision was defined as the proportion of identified preys that matched the benchmarking list; recall reflected the proportion of true benchmarking proteins recovered. Enzyme-specific PR curves were generated by averaging precision and recall across all baits in each enzyme group. Custom Python script Scikit-learn was used to generate the PR curves^80^. In the second approach (Supplementary Fig. 2b), to allow for a more holistic view of the dataset, proximal interactors were ranked by SAINT scores in intervals of 50 per bin and precision and recall were calculated over a full range of SAINT score thresholds, from 0 to 1. The Area Under Curve (AUC) was calculated for each enzyme at recall 0.5^81^.

### GO enrichment dot plot

A GO enrichment dot plot was generated using in-house developed code in Python. The analysis was performed via g:Profiler, and the resulting .csv files (containing adjusted p-values and gene counts for each GO term) were used as input for the Python visualization pipeline. The top 3 and 5 GO:CC terms (by lowest p-value) per sample were displayed for HeLa and HEK293 datasets respectively, and the data were filtered for GO term size ≤ 1500. In these plots, the square color represented the adjusted P-value for enrichment, while the square size corresponded to the number of preys contributing to each GO term.

### Data visualization

Bait-prey dot plots and bait-versus-bait comparisons were visualized using ProHits-viz^82^ [http://prohits-viz.org]. In the dot plot tool, any prey passing the selected BFDR threshold (≤ 1%) for at least one bait had its spectral count values displayed across all baits. The node edge color represented the BFDR of the prey for each bait, while the node’s color gradient indicated the corresponding spectral count values. Node size reflected the relative prey counts across all baits or conditions, normalized to the bait with the highest spectral count.

The UpSet plots were generated using the UpsetR package^83^, bar plots, dot plots, violin plots and heatmaps were generated using in-house Python scripts (available upon request). Log_2_-transformed average spectral counts across replicates for preys passing SAINT BFDR ≤ %1 in at least one condition (i.e., for at least one enzyme and biotinylation timepoint) were clustered by Pearson correlation and average linkage using the Python seaborn package’s clustermap function^84^. Violin plots and figures were finalized and formatted for publication using Adobe Illustrator (version 28).

### Data availability

Data has been deposited as a complete submission to the MassIVE repository (https://massive.ucsd.edu/ProteoSAFe/static/massive.jsp) in four datasets and assigned the accession numbers MSV MSV000098673 (dataset 1 – ten biotin ligases fused to LMNA at two different time windows, no biotin and 6-hour biotinylation), MSV000098677 (dataset 2 – four biotin ligases fused to LMNA at four time windows, no biotin, 5 minutes, 15 minutes, and 1 hour), MSV000098681 (dataset 3 – three biotin ligases fused to six different baits representing diverse subcellular compartments in HEK293 cells at 15 minutes of biotinylation), and MSV000098669 (dataset 4 – two biotin ligases fused to eight cytoskeletal and endosomal baits in HeLa cells at 15 minutes of biotinylation.) The ProteomeXchange accession is PXD066731, PXD066734, PXD066742 and PXD066728 respectively.

## Supplementary information

The manuscript is accompanied by 4 supplementary Figures (this document) and 10 Supplementary Tables, presented as Excel files.

## List of tables

Supplementary Table 1: List of constructs used in this study

Supplementary Table 2: Protein identified in the eGFP negative controls in normal growth conditions and at the 6 h time point post biotin supplementation

Supplementary Table 3: SAINTexpress results and reproducibility metrics

Supplementary Table 4: Prey overlap across 6 h biotin and no-added-biotin conditions (supports Fig. 2a)

Supplementary Table 5: BioGRID overlap for LMNA at 6h (supports Fig. 2d)

Supplementary Table 6: True/False positive (GO:CC) for LMNA at 6h (supports Fig. 2e)

Supplementary Table 7: Enrichment of nuclear lumen preys at short time points

Supplementary Table 8: GO CC enrichment for Dataset 3

Supplementary Table 9: GO CC enrichment for Dataset 4

Supplementary Table 10: Immunofluorescence co-localization analysis

**Sup. Fig 1.**
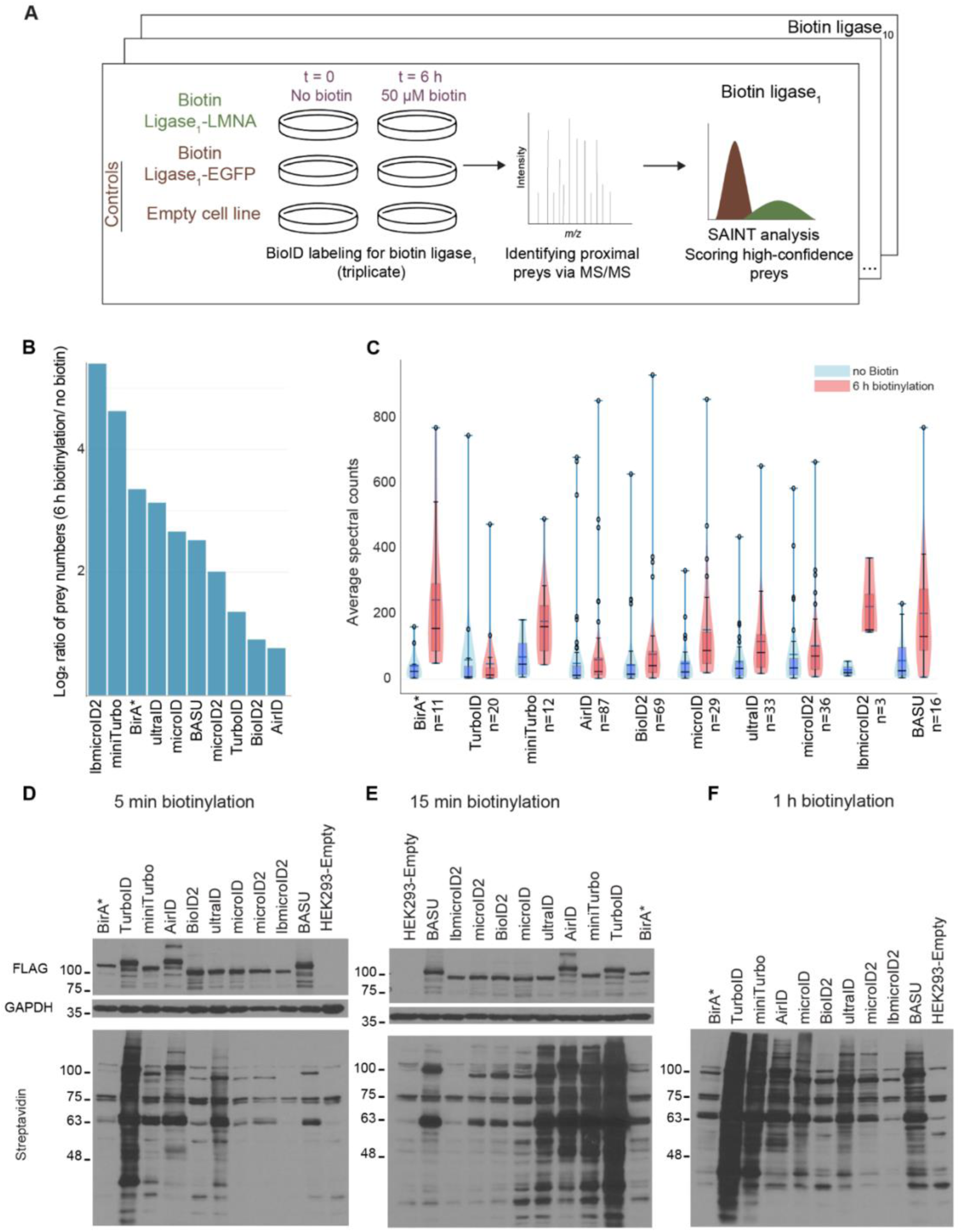
Experimental workflow and comparative evaluation of ten biotin ligases using LMNA. **A.** Schematic overview of the experimental design. HEK293 Flp-In T-REx cell pools inducibly expressing LMNA fused to each of ten biotin ligases were incubated with or without 50 µM biotin for 6 h. Each condition was performed in biological triplicates and compared against biotin ligase–EGFP controls and empty cells. Proximal proteins were identified by mass spectrometry, and high-confidence interactors were scored using SAINTexpress. **B**. Violin plots showing the distribution of mean spectral counts for prey proteins detected in both the no-added-biotin (blue) and 6-h biotinylation (red) conditions across all tested ligases. The number of shared preys is indicated (n = x). **C–E.** Western blot analysis of HEK293 Flp-In T-REx cells expressing LMNA–biotin ligase fusions following 5 min (C), 15 min (D), or 1 h (E) of biotin labeling. FLAG blots indicate bait expression, streptavidin-HRP shows biotinylation levels, and GAPDH serves as a loading control.

**Sup. Fig 2.**
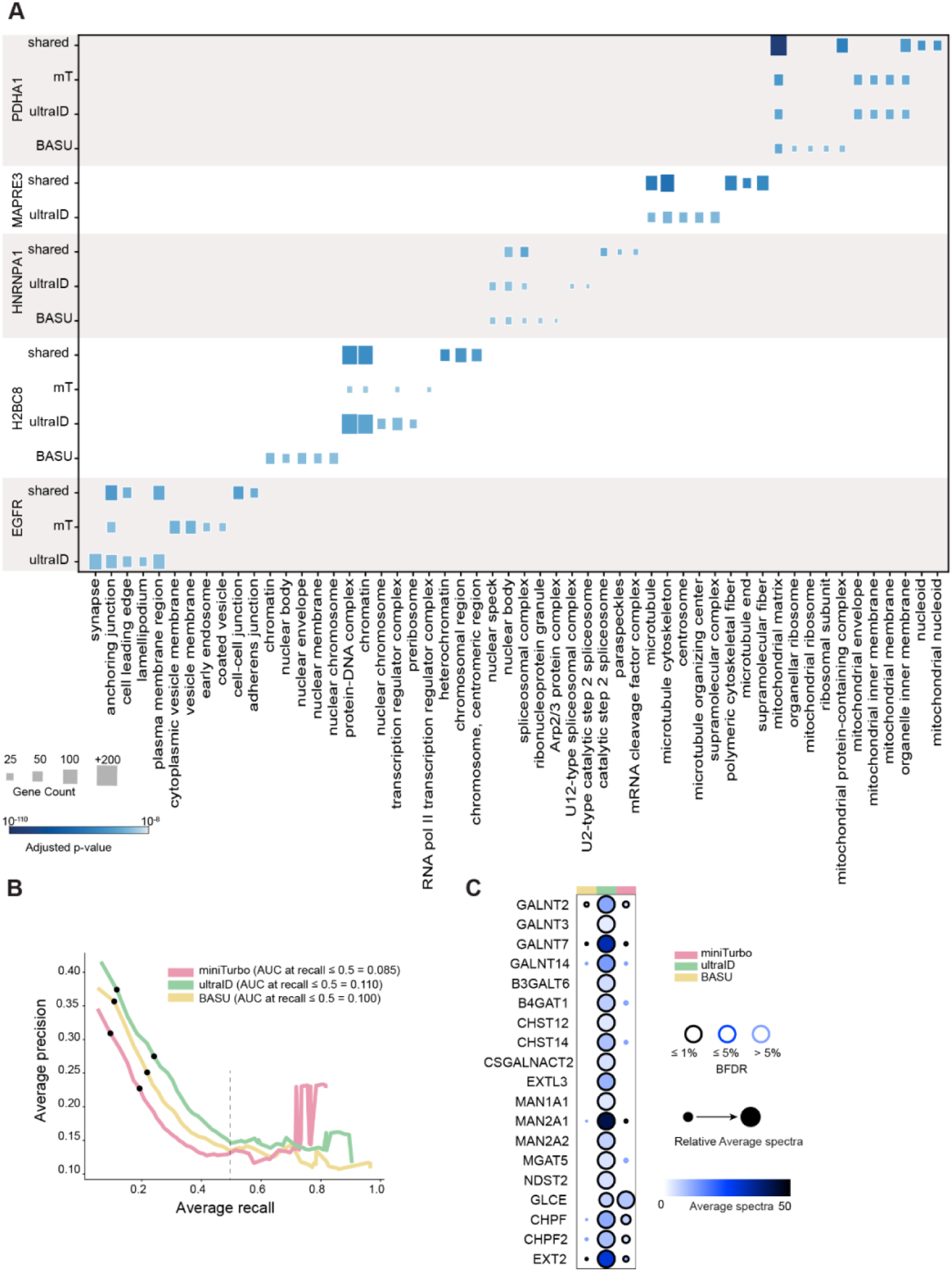
Validation of bait expression and GO:CC enrichment of high-confidence interactors in HeLa cells. **A.** GO cellular component (GO:CC) enrichment analysis of high-confidence proximity interactors for each bait fused to miniTurbo, BASU or ultraID. The top 5 enriched GO:CC terms per bait are shown (based on adjusted p-value), filtered for terms with ≤1500 genes. Square size reflects gene count, and color scale indicates adjusted p-values. **B.** Precision-recall (PR) curves evaluating the performance of each biotin ligase based on SAINT score. Ground-truth interactors were defined based on Gene Ontology Cellular Component (GO:CC) terms relevant to each bait (see Methods). Preys were ranked by SAINT score and grouped into intervals of 50. Average PR curves are shown for each biotin ligase, aggregated across all baits. AUC: Area Under Curve. The dashed line indicates recall at 0.5. On each curve, the top dot indicates a SAINT score of 0.95, and the bottom dot indicates a SAINT score of 0.7. **C.** Dot plot of selected Type II Golgi membrane proteins enriched by ST6GALNAC1 fusions.

**Sup. Fig 3.**
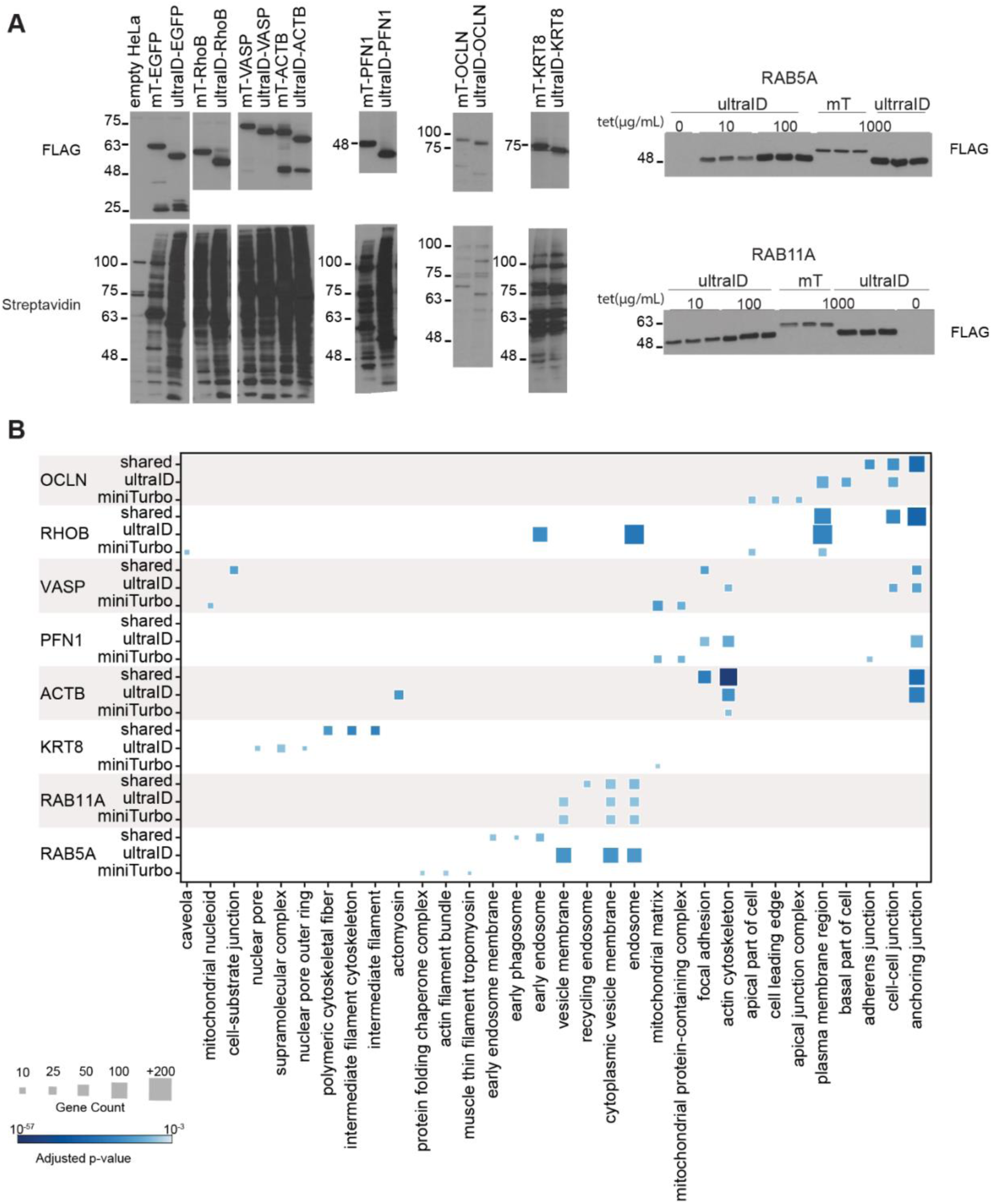
Validation of bait expression and GO:CC enrichment of high-confidence interactors in HeLa cells. **A.** Western (FLAG) and Far-Western (streptavidin) blot analysis of FLAG-tagged bait–enzyme fusion expression and associated biotinylation in HeLa cells. For RAB5A and RAB11A fusions, expression of ultraID constructs was tuned using lower tetracycline concentrations (10 µg/mL) to match levels across enzymes. **B.** GO cellular component (GO:CC) enrichment analysis of high-confidence proximity interactors for each bait fused to miniTurbo or ultraID. The top three enriched GO:CC terms per bait are shown (based on adjusted p-value), filtered for terms with ≤1500 genes. Square size reflects gene count, and color scale indicates adjusted p-values.

**Sup. Fig. 4.**
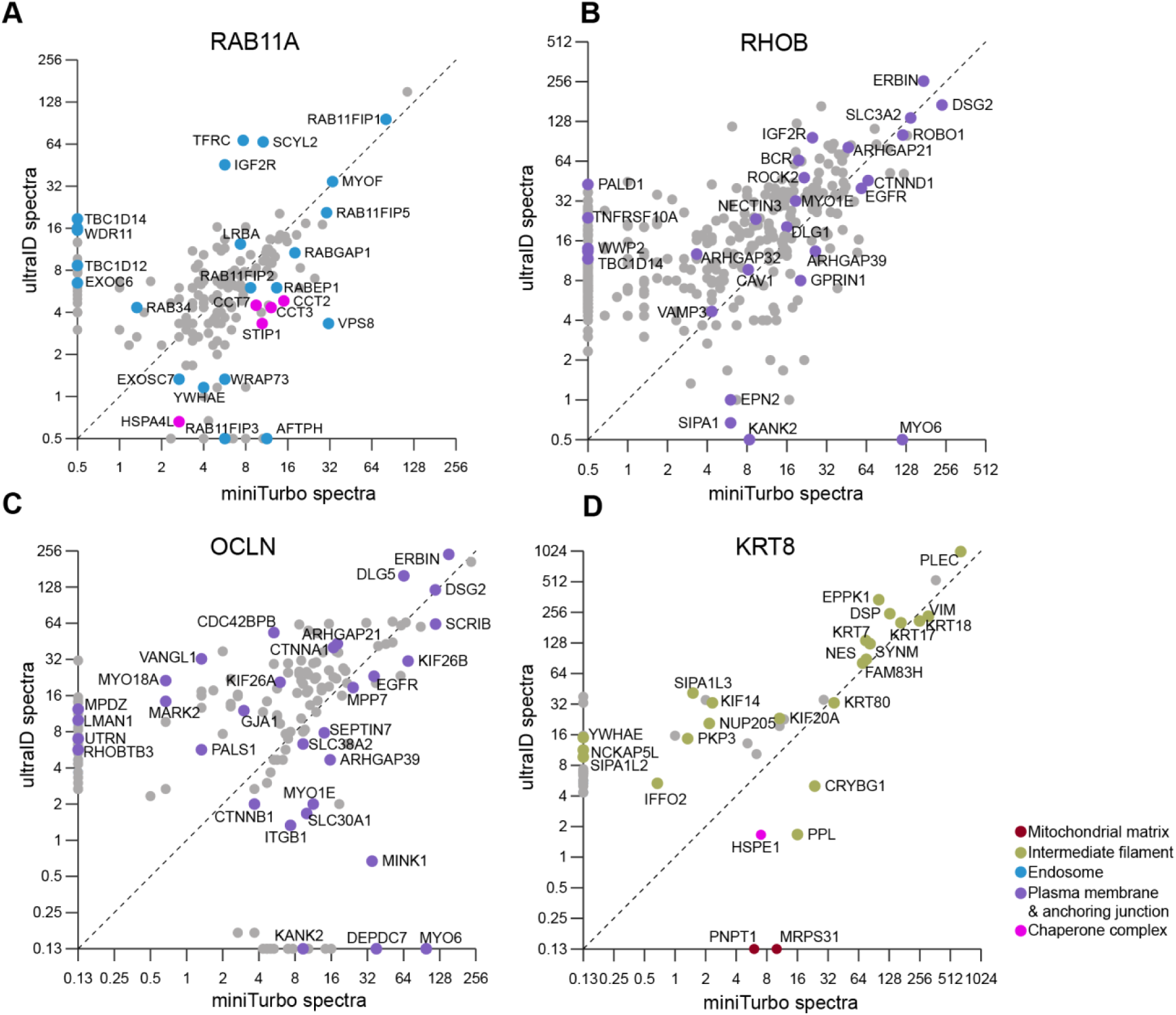
ultraID consistently recovers higher-abundance proximity interactors across baits. **A**–**D**. Scatter plots comparing abundance (log₂-transformed average spectral counts) of high-confidence proximity interactors for RAB11A (A), RHOB (B), OCLN (C), and KRT8 (D) when fused to miniTurbo (x-axis) or ultraID (y-axis). Each point represents a prey protein. The dashed diagonal indicates equal abundance between enzymes. Preys are colored by GO:CC annotation: mitochondrial matrix (red), intermediate filament (olive), endosome (cyan), actin plasma membrane and anchoring junction (purple), and chaperone complex (magenta).

## Notes

### Competing Interest Statement

The authors have declared no competing interest.

